# Gene Family Complexity and Expression Divergence as a Mechanism of Adaptation In Coral

**DOI:** 10.1101/2021.04.28.441826

**Authors:** Bradford Dimos, Madison Emery, Nicholas MacKnight, Marilyn Brandt, Jeffery Demuth, Laura Mydlarz

**Affiliations:** Department of Biology, University of Texas at Arlington, Arlington, TX 76019, USA; Center for Marine and Environmental Studies, University of the Virgin Islands, St. Thomas, US Virgin Islands 00802, USA

## Abstract

Gene family complexity and its influence on expression dynamics has long been theorized to be an important source of adaptation in natural systems through providing novel genetic material and influencing gene dosage. There is now growing empirical support for this theory; however, this process has only been demonstrated in a limited number of systems typically using recently diverged species or populations. In particular, examples of how this process operates in basal animals with deeper species splits has not been well explored. To address this issue, we investigated the evolution of gene family complexity in five species of common Caribbean coral. We demonstrate widespread divergence in gene repertoires owing to slow rates of gene turnover occurring along deep species splits. The resulting differences in gene family complexity involve numerous biologic processes, shedding light on to the selective forces that have influenced the evolution of each species. By coupling these findings with gene expression data, we show that increased gene family complexity promotes increased expression divergence between species, indicating an interplay between gene family complexity and expression divergence. Finally, we show that immune genes are evolving particularly fast demonstrating the importance of interactions with other organisms in the evolutionary history of Caribbean corals. Overall, these findings provide support for gene copy number change as an important evolutionary force in Caribbean corals, which may influence their ability to persist in a rapidly changing environment.

## Intro

Understanding the genetic mechanisms which facilitate how organisms generate phenotypes which are adapted to their environment is a major goal of modern biology. This process is commonly investigated through quantifying gene expression patterns, as changes in gene expression can operate over short time scales to promote acclimation (Bay & Palumbi, 2015; Kenkel & Matz, 2017). Expression patterns, however, can be heritable across generations (Monks et al., 2004; Price et al., 2011), and facilitate adaptation at the population or species level (Fraser, 2011; Whitehead & Crawford, 2006). How an organism expresses its genes to match its environment is influenced by the underlying gene repertoire, as gene specificity influences under what circumstances a gene will be expressed. This is especially true in mutli-copy gene families which evolve through gene gain and loss events which change the amount of genetic material available for transcription (Meyers et al., 2004). The resulting gene copy number variation can facilitate adaptation by increasing genetic novelty (Deng, Cheng, Ye, He, & Chen, 2010; Hirase, Ozaki, & Iwasaki, 2014) and promoting the diversification of organismal traits (Chain et al., 2014; Tejada-Martinez, de Magalhães, & Opazo, 2021), through the processes of gene neofunctionalization and subfunctionalization (Force et al., 1999; Ohno, Susumu, 1970). However, the role of the resulting transcriptional complexity generated by gene family evolution has received relatively little attention as a mechanism of adaptation.

In addition to providing genetic novelty, gene copy number directly influences expression patterns, as a transcriptionally active gene duplicate will serve to increase expression (Kondrashov, 2012; Schuster-Böckler, Conrad, & Bateman, 2010). Such changes in gene dosage if advantageous will fix (Brown, Todd, & Rosenzweig, 1998). Alternatively deleterious changes in gene dosage may be resolved through epigenetic silencing (K. M. Huang & Chain, 2021) or sub-division of the two gene copies (De Smet, Sabaghian, Li, Saeys, & Van de Peer, 2017). The interaction of both transcriptional complexity and gene expression affects how an organism utilizes its underlying genetic material, as gene copy number dictates the transcriptional complexity and dosage of a gene family. Despite a growing realization of the importance of gene copy number variation and expression divergence in adaptation (Gilad, Oshlack, & Rifkin, 2006; Yun Huang et al., 2019) these processes are rarely investigated in tandem.

Reef-building corals are a good system in which to study how transcriptional complexity mediated by gene family evolution contributes to adaptation. Adult corals are sessile meaning that they cannot track favorable environmental conditions and instead must be able to dynamically adjust gene expression profiles in order to persist during unfavorable conditions. Increased transcriptional complexity through gene copy number changes may be a way to accommodate this sessile life-history strategy as seen in plants (Prunier, Caron, & MacKay, 2017; Würschum, Boeven, Langer, Longin, & Leiser, 2015). Additionally, corals lack differentiated organs and have relatively homogenous tissue limiting the influence of tissue-specific expression patterns. Demographically, corals have historically possessed large effective population sizes (Matz, Treml, Aglyamova, & Bay, 2018; Prada et al., 2016) and high genetic connectivity (S. W. Davies, Treml, Kenkel, & Matz, 2015), conditions which favor efficient selection. Under such scenarios large scale mutations, such as changes in copy number of a gene, are expected to rise to fixation or be purged from the population quickly. Thus, variation in transcriptional complex may be an important mechanism of species-level adaptation in reef-building corals. In support of this, some investigations have reported variation in multi-copy gene families between coral species (Dimos, Butler, Ricci, MacKnight, & Mydlarz, 2019; Hamada et al., 2013; Shinzato et al., 2020), however a systematic evaluation of this process between species from different genera has not been performed.

While historical environmental conditions allowed corals to thrive, current environmental conditions are highly variable and are causing widespread coral loss across the globe (Maynard et al., 2015; Stuart-Smith, Brown, Ceccarelli, & Edgar, 2018). In particular, Caribbean coral reefs have been severely depleted (Gardner, 2003), owing to the combined effects of thermally induced coral bleaching and the emergence of diseases effecting reef-building corals (Smith et al., 2013; van Woesik & Randall, 2017). This underscores the differences between the historical selective forces that shaped coral communities such as benthic space and light availability (Alvarez-Filip, Dulvy, Gill, Côté, & Watkinson, 2009) versus the current selective forces acting to reshape these ecosystems including marine heatwaves and coral diseases (Aronson & Precht, 2001; Baker, Glynn, & Riegl, 2008; Precht, Gintert, Robbart, Fura, & van Woesik, 2016; Walton, Hayes, & Gilliam, 2018). Thus, the environmental conditions facing corals today are different than the conditions these organisms evolved under, meaning corals must either adapt or face extirpation (Putnam, Barott, Ainsworth, & Gates, 2017).

Most studies investigating the adaptive capacity of corals utilize intraspecies data by focusing on processes such as gene expression plasticity (Kenkel & Matz, 2017), or single-nucleotide variants (Bay & Palumbi, 2014; Fuller et al., 2020). However, interspecies variation in gene repertoires has received relatively little attention particularly in the context of evolutionary adaptation. In this investigation we use gene expression data to investigate the evolution of transcriptional complexity, defined as changes in the number of predicted sequences in orthologous gene families, in five ecologically important species of Caribbean corals: *Orbicella faveolata*, *Montastraea cavernosa*, *Colpophyllia* natans, *Porites astreoides*, and *Siderastrea siderea*. Our analysis demonstrates wide-spread variation in both gene repertoire content and transcriptional complexity in gene families involved in numerous biological processes. We expand upon these results to show that transcriptional complexity and species-level adaptive shifts in expression are linked. Finally, we demonstrate that immune related genes have particularly high rates of both transcriptional complexity and expression divergence highlighting the role of immune evolution in the history of these species.

## Methods

### Coral collection

Five colonies of *O. faveolata*, *M. cavernosa*, *C.* natans, *P. astreoides*, and *S. siderea* were collected via SCUBA from Brewer’s Bay St. Thomas (18.34403, −64.98435) in June 2017 under the Indigenous Species Research and Export Permit number CZM17010T. Colonies were transported to the Center for Marine and Environmental Studies at the University of the Virgin Islands and allowed to acclimate for two weeks in large flow-through holding tanks fed with filtered sea water from the bay which received natural sunlight under a neutral density shade cloth. After this acclimation period samples were flash frozen in liquid nitrogen for transport back to the University of Texas Arlington.

### RNA extraction and sequencing

Total RNA was extracted from coral fragments using the Ambion RNAeasy kit with DNase according to manufactures protocol and eluted in 100μl of elution buffer. RNA integrity and concentration were checked with an Agilent Bioanalyzer 2100 using the RNA 6000 Nano kit, and all samples with a RIN greater than 7.0 and above 1 μg of total RNA were used for sequencing. Samples were sequenced through the Novagene company and mRNA libraries were prepared with the Illumina TruSeq RNA library prep kit which uses Poly-A tail enrichment to purify mRNA. After library prep samples were sequenced on an Illumina HiSeqX at Novagene company. Five colonies of each *M. cavernosa* and *C. natans* were sequenced, while four colonies of the other species were sequenced. Raw reads from this experiment can be found on the NCBI short read archive (PRJNA723585).

### Transcriptome assembly and cleaning

Transcriptome assemblies were generated with Trinity (Grabherr et al., 2011) after removing adapters and low-quality reads with trimmomatic (Bolger et al., 2014). For *O. faveolata* a genome is available on NCBI and we chose to use this assembly (assembly ofav_dov_v1) (Prada et al., 2016). For *M. cavernosa* a draft genome assembly is available (https://matzlab.weebly.com/data--code.html) and we chose to use this genome assembly to generate a genome-guided transcriptome assembly for *M. cavernosa*. Reads from our five *M. cavernosa* colonies were mapped to the draft genome with the read mapping program tophat2 (Kim et al., 2013) using default parameters. The merged bam files were then used in the Trinity genome-guided approach to generate our transcriptome. For *C. natans, P. astreoides,* and *S. siderea* we used the filtered reads to generate de novo transcriptome assemblies in Trinity.

### Symbiont filtration

As transcriptome assemblies derived from adult corals contain transcripts arising from symbiont containments these must be filtered out prior to downstream analysis. To filter out symbiont transcripts we utilized a previously described approach to symbiont filtration (Sarah W. Davies, Marchetti, Ries, & Castillo, 2016). In short, multiple coral assemblies (Davies, Marchetti, Ries, & Castillo, 2016; Kirk, Howells, Abrego, Burt, & Meyer, 2017; Moya et al., 2012; van de Water et al., 2018) were combined to create a coral database, while assemblies from multiple genera of symbionts (Aranda et al., 2016; Liu et al., 2018; Parkinson et al., 2016; Shoguchi et al., 2021) were combined to create a symbiont database. Transcripts from each of our coral assemblies were then queried using blastn against each database and a transcript was retained only if it had greater than 80% identity over at least 100bp to the coral database and that transcript did not have a lower evalue when blasted against the symbiont database. This was carried out on the *C. natans*, *P. astreoides* and *S. siderea* transcriptome assemblies. For *O. faveolata* and *M. cavernosa* no symbiont filtration was performed as the process of reference mapping to create transcriptomes eliminates symbiont containments.

### Transcript Filtering

To interpret transcriptome complexity in terms of gene family evolution it is necessary to have high confidence in the breadth of the transcripts sampled relative to the gene content, and that there is a one-to-one correspondence between transcripts and genes. We took several steps to ensure these relationships hold before conducting subsequent analyses. First, each assembly was filtered for longest isoform to attempt to remove extraneous transcripts in our assemblies which derive from splice-variants of a gene using the get longest isoform script available through trinity (Grabherr et al., 2011). Next the program TransDecoder (Haas et al., 2013) was used to extract the longest open reading frame of each transcript and this reading frame was utilized to generate a predicted peptide sequence in TransDecoder. This step was performed for each species with the exception of *O. faveolata* for which we utilized the proteome available from NCBI. The predicted proteomes for all species were then collapsed for similar sequences using the program cd-hit (Huang et al,. 2010) at a similarity level of 0.9 to further exclude transcripts which are likely derived from alternative splice-variants of a gene. For the two species with available genomes we mapped the predicted peptide sequences back to the genome assembly using tblastn to identify transcripts which shared a mapping start site and are likely isoforms. We removed these likely isoforms by retaining only the longest transcript with an overlapping start site in the assembly. To ascertain the quality of these assemblies after transcript filtration we employed the program Benchmarking Universal Single Copy Orthologs (BUSCO) (Simão et al., 2015), and checked the completeness of each assembly against the core metazoan database. To generate gene ontology annotations from our assemblies we utilized the annotation software EggNog mapper (Huerta-Cepas et al., 2017, 2019) through the online web portal against the eukaryote database. To annotate individual contigs we used blastp against the swiss-prot uniprot database (The UniProt Consortium et al., 2021).

### Orthogroup assignment

To identify homologous genes across our species we utilized the program Orthofinder2 (David M. Emms & Kelly, 2015, 2019). This software groups transcripts into groups containing both orthologs and paralogs and these groups will here out be referred to as gene families. A species tree was generated through the program species tree from all genes (STAG) (D.M. Emms & Kelly, 2018) as implemented within Orthofinder2 where branch lengths are measured in amino acid substitutions. We then converted the calculated branch lengths into time based on the estimated divergence times between species (Pinzón C., Beach-Letendre, Weil, & Mydlarz, 2014).

### Gene family evolution

To quantify the rate of gene family evolution we employed the program Café3 (Han, Thomas, Lugo-Martinez, & Hahn, 2013). This program utilizes an ultrametric species tree with a set of gene families to determine the background rate of gene gain and loss per gene per unit time across the provided species tree. This rate of gene turnover is then used a null model to test for gene families which are experiencing significant deviations from this rate. For our analysis the species tree generated through STAG was converted into an ultrametric tree using phytools (Revell, 2012) with the option “extend”. The list of gene families was refined to contain only families which contained at least one transcript in all species to improve our estimate of gene turnover which encompassed 6324 gene families. To account for the role that errors in our assemblies may have played in over-inflating our estimate of gene turnover we utilized the error model script included in the Café3 package (Han et al., 2013). This script iteratively searches *a priori* defined error distributions to identify which distribution maximizes the probability of observing the data. The error model with the highest probability of observation was then used to calculate gene turnover. To determine if a single global rate of turnover or multiple rates should be utilized on our dataset, we performed 100 simulations with our geneset to generate a null distribution of log-likelihood ratios comparing a tree-wide rate of turnover to two rates of turnover where the species with the highest number of gene families undergoing rapid evolution (*M. cavernosa*) was given a species-specific rate. The observed likelihood ratio between a single versus multiple rates of turnover was compared to the expected distribution to generate a p-value. To account for the possibility that assembly quality may have influenced the estimates of gene family evolution we constructed a multiple linear model to regress transcript number, number of present BUSCOs and number of duplicated BUSCOs against the number of rapidly evolving gene families in each species.

### Gene ontology enrichment

To understand the processes which are undergoing rapid evolution in these species we utilized the R script Gene Ontology (GO) with Mann-Whitney U test (GO_MWU) (Wright, Aglyamova, Meyer, & Matz, 2015), to perform a fisher’s exact test where contigs in rapidly evolving gene families were assigned a value of 1 while all other contigs were assigned a value of 0. The enrichments were run against the background of all of the peptide sequences present within the assemblies. This was performed in order to identify enrichments within the background of each species rather than testing for enrichments based only within conserved genes, and thus represents a conservative statistical approach. This method was employed separately for the transcripts in gene families with increased complexity as well as the transcripts in gene families with decreased complexity seperately. For each species identical parameters were used for biological process (BP) with the category smallest set to 50, largest to 0.1 and clusterCutHeight at 0.25.

### Expression divergence

In order to determine which genes demonstrate species-specific expression shifts we mapped reads from each coral colony to a holobiont transcriptome with the read mapping program salmon (Patro, Duggal, Love, Irizarry, & Kingsford, 2017) using an index size of 31 in each species. The holobiont transcriptome was generated by combining the host transcripts with available transcriptomes from representatives of four genera of Symbiodiniaceae: *Symbiodinium microadriaticum* (Aranda et al., 2016), *Breviolum minutum* (Parkinson et al., 2016), *Cladocopium goreaui* (Liu et al., 2018), *Durusdinium trenchii*, (Shoguchi et al., 2021). Read counts belonging to the host were then extracted and summarized to gene family level with the R package TXimport (Soneson, Love, & Robinson, 2015) using the “salmon” option. These expression profiles were then combined across species, filtered for low abundance transcripts (<10 average counts) and rlog normalized by species in the R package DESeq2 (Love, Huber, & Anders, 2014). Species-level expression shifts were quantified using the R package EVE using a β shared test (Rohlfs & Nielsen, 2015). Where the rlog normalized counts of each gene family as well our generated phylogenetic tree were used as input files. This program identifies gene families with both high levels of between species expression variance represented by low β values versus gene families with high levels of within species expression variance represented by high β values. Low β values indicate divergent shifts in expression, while high β values indicate genes with evolutionarily conserved expression profiles. β values were -log10 transformed to generate our expression divergence metric. To determine if rapidly evolving gene families possess higher expression divergence than single copy gene families, we compared the number the number of gene families with significant expression divergence (p<0.01) contained within the rapidly evolving gene families to the single copy gene families using a fisher’s exact test. To account for the influence of gene family size on expression divergence we regressed the number of transcripts in a gene family against the expression divergence value of the gene family. To compare the evolutionary patterns of immune genes, the gene families with the annotation GO:0002376 “Immune System Process” were extracted and the expression divergence was compared between the rapidly evolving immune gene families, the remaining rapidly evolving gene families and the single copy gene families with an anova with Tukeys HSD test for multiple comparisons. All statistical analysis were carried out in the R programing language (*R Core Team (2020)*).

## Results

### Assembly statistics and Orthogroup Assignement

With our filtering strategy we produced assemblies containing 23378 peptide sequences for *O. faveolata*, 34315 for *M. cavernosa*, 52753 for *C. natans*, 39641 for *P. astreoides* and 19759 for *S. siderea*. The percentage of BUSCOs present in these assemblies ranged from 83-96% and the number of duplicated BUSCOs ranged from 1.8% to 3.1%. Collectively we identified 27209 gene families. This resulted in 6324 gene families which had at least one transcript in all species, of which 2009 are single copy across all species. By considering only transcripts assigned to gene families containing at least one transcript per species we further refined our assemblies to 9322 transcripts in *O. faveolata*, 11347 in *M. cavernosa*, 10919 in *C. natans*, 10767 in *P. astreoides* and 9185 in *S. siderea* (Figure 1a). The distribution of the number of transcripts in gene family size class within this refined list is similar across species (Figure 1a); most transcripts are in gene families with a single member and the number of transcripts in each gene family size class decreases as size class increases. Our phylogenetic tree based on these gene families recapitulates the known phylogenetic relationship between these species of coral (Figure 1b).

**Figure 1:**
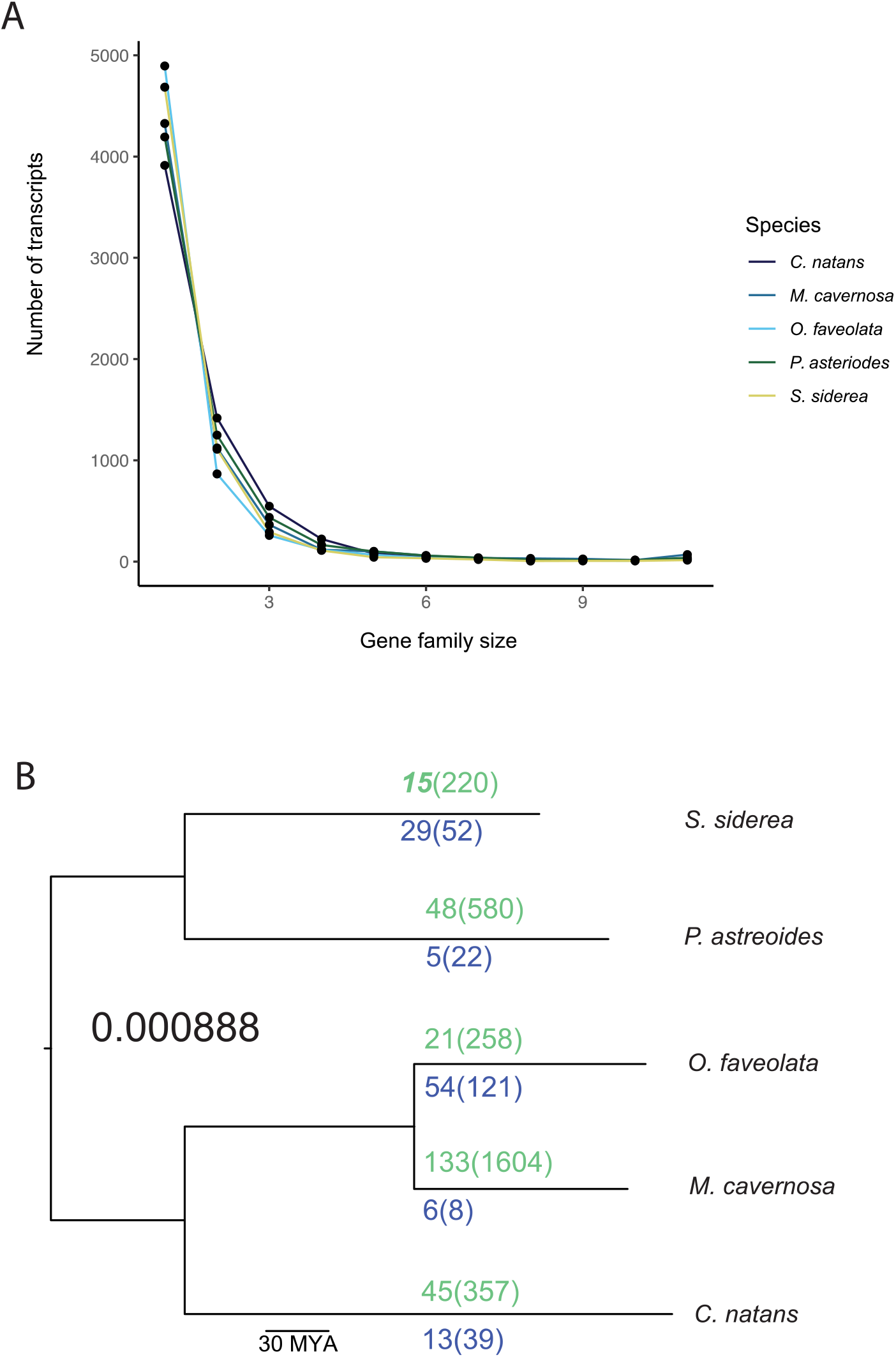
Quantifying gene family evolution. A) Shown is the distribution of the number of transcripts contained in each size of gene family where each species is represented by a different color. B) Species tree shown with the background rate of gene turnover. The number of rapidly evolving gene families in each species is shown on the branches where the number of gene families with increased complexity are listed in green and the number of gene families with decreased complexity are listed in blue. The number of transcripts contained with these gene families are shown inside the parenthesis.

### Orthogroup evolution

The best fitting error model to our data was 1.22e-5, which was used in our calculation of gene turnover. By evaluating transcriptome complexity across our focal species, we were able to identify a background gene gain and loss per gene per million years (MYA) of 8.88e-4 (Figure 1b). We identified 352 gene families that have experienced rapid evolution (p<0.01) and are considered to have evolved changes in complexity. The number of rapidly evolving gene families in a species was not influenced by assembly quality as the model quantifying the association between number of transcripts, number of BUSCOS present and number of duplicated BUSCOS was not significant (p=0.137). Additionally, when regressed individually none of the three parameters were significantly associated with the number of rapidly evolving gene families (p>0.05). The number of rapidly evolving gene families varied by species: 75 in *O. faveolata*, 139 in M*. cavernosa*, 45 in *C. natans*, 53 in *P. astreoides* and 44 in *S. siderea* (Figure 1b). Despite having the highest number of rapidly evolving gene families, including a separate rate of evolution for *M. cavernosa* was not statistically supported as the observed likelihood ratio between a global compared to multiple rates of gene turnover outperformed all 100 simulations (p<0.01) (Figure S.1), supporting a single tree-wide rate of turnover.

*O. faveolata* had 75 gene families undergoing rapid evolution, of which 21 had increased complexity and 54 had decreased complexity. The gene families with increased complexity contained a total of 258 transcripts of which 245 (95%) were annotated while the gene families with decreased complexity contained 121 transcripts of which 108 (89%) were annotated (Figure 1b). The transcripts in gene families with increased complexity were enriched for 16 GO terms (p<0.05) (Table S.1), while the transcripts in gene families with decreased complexity were enriched for five terms (Table S.2).

*M. cavernosa* had 139 gene families undergoing rapid evolution, of which 133 had increased complexity while six had decreased complexity. The transcripts with increased complexity contained 1604 transcripts of which 1155 (72%) were annotated while the gene families with decreased complexity contained eight transcripts of which four (50%) were annotated (Figure 1b). The gene families with increased complexity were enriched for 64 GO terms (Table S.3), while the gene families with decreased complexity did not possess any GO enrichments.

*C. natans* had 45 gene families undergoing rapid evolving of which 32 had increased complexity and 13 of which had decreased complexity. The gene families with increased complexity contained 357 transcripts of which 197 were annotated (55%) while the gene families with decreased complexity contained 39 transcripts of which 14 (36%) were annotated (Figure 1b). The gene families with increased complexity were enriched for 78 GO terms (Table S.4). The gene families which had reduced complexity in this species were enriched for 12 GO terms (Table S.5).

In *P. astreoides* there were 53 gene families undergoing rapid evolution of which 48 had increased complexity while five had decreased complexity. The transcripts in gene families with increased complexity contained 580 transcripts of which 493 (85%) were annotated while the gene families with decreased complexity contained 22 transcripts of which all were annotated (Figure 1b). The gene families with increased complexity were enriched for 29 GO terms (Table S.6), while the transcripts in gene families with decreased complexity did not have any GO enrichments.

In *S. siderea* 44 gene families were undergoing rapid evolution of which 15 had increased complexity and 29 had decreased complexity. The gene families with increased complexity contained 220 transcripts of which 214 (97%) were annotated. There were 52 transcripts in gene families which had decreased complexity of which 47 (90%) were annotated (Figure 1b). The transcripts in gene families with increased complexity were enriched for 16 GO terms (Table S.7), while the transcripts in gene families with decreased complexity did not possess any GO enrichments.

Across the species there were a total of 41 enriched GO terms which fell under the parent terms “Immune System Process” GO:0002376 or “Metabolic process” (GO:0008152). Of these enriched terms all had increased complexity, except for two terms in *O. faveolata* which demonstrated decreased complexity (Figure 2).

**Figure 2:**
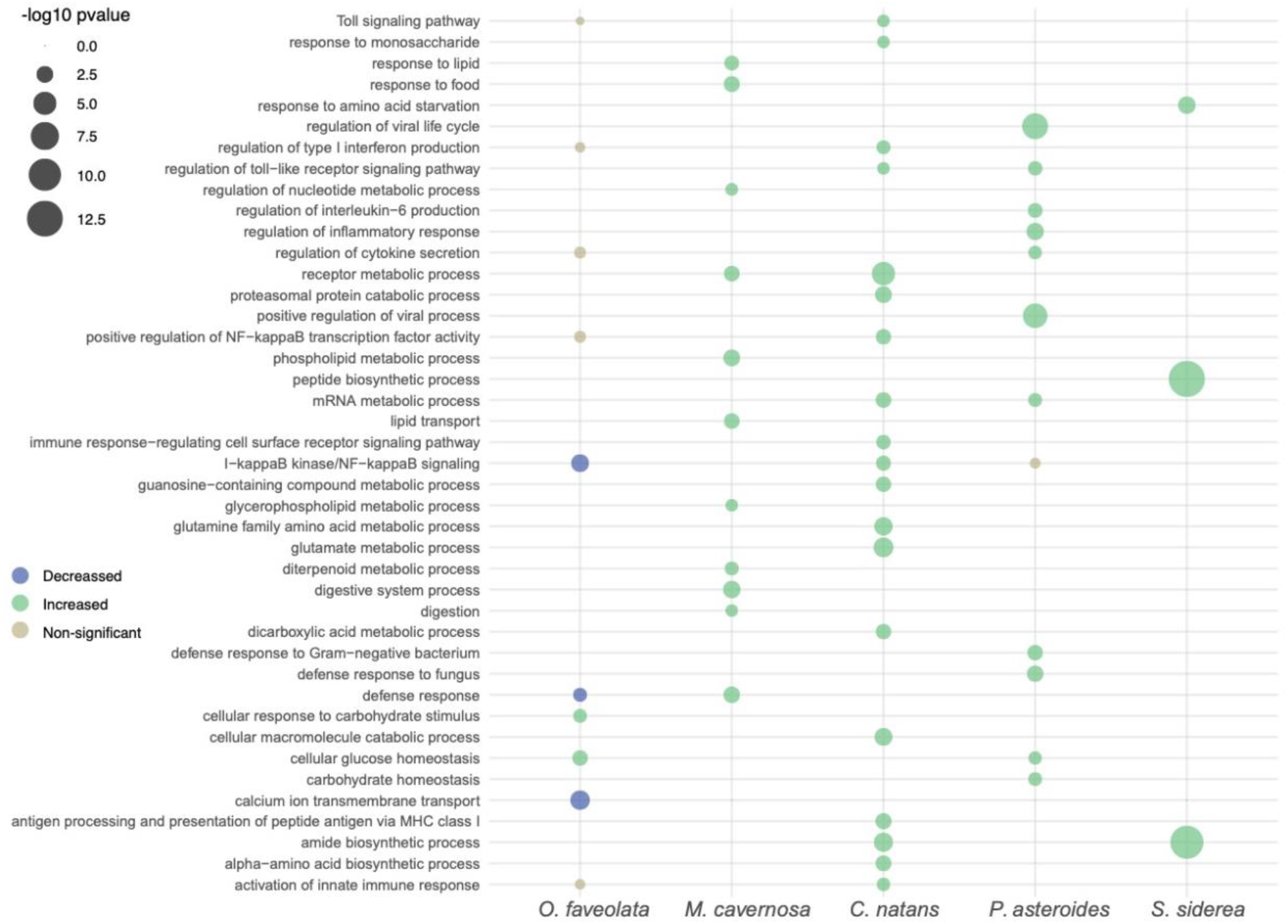
GO enrichments for immune and metabolic gene families. Shown is a bubble plot of GO terms under the parent term “Immune System Process” GO:0002376 or “Metabolic process” (GO:0008152). Size of the circle corresponds to the significance (-log10 transformed p-value) after false discovery rate correction, and color denotes if the term enrichment is in gene families which are expanding (green) or contracting (blue) or if the term is non-significant after FDR correction (grey).

### Expression divergence

To test if the gene families with evolved changes in complexity demonstrated species-specific expression shifts, we quantified gene expression patterns across the gene families. Of the 6324 gene families which were present across all species 5451 were highly expressed including 313 of the 352 rapidly evolving gene families. Overall, 155 gene families demonstrate higher expression divergence between species than expression diversity within species (p<0.01) (Figure 3) and are considered to have divergent expression patterns. 4.8% of the rapidly evolving gene families were contained within the gene families with divergent expression patterns, while only 2.5% of single copy gene families also displayed divergent expression. Overall, rapidly evolving gene families were more likely to demonstrate expression divergence (Odds ratio = 1.93, p<0.05, fisher’s exact test) (Figure 3). There was a significant association between gene family size and expression divergence (p<0.01), however gene family size explains very little of the variance in expression divergence (R^2 = 0.0027).

**Figure 3:**
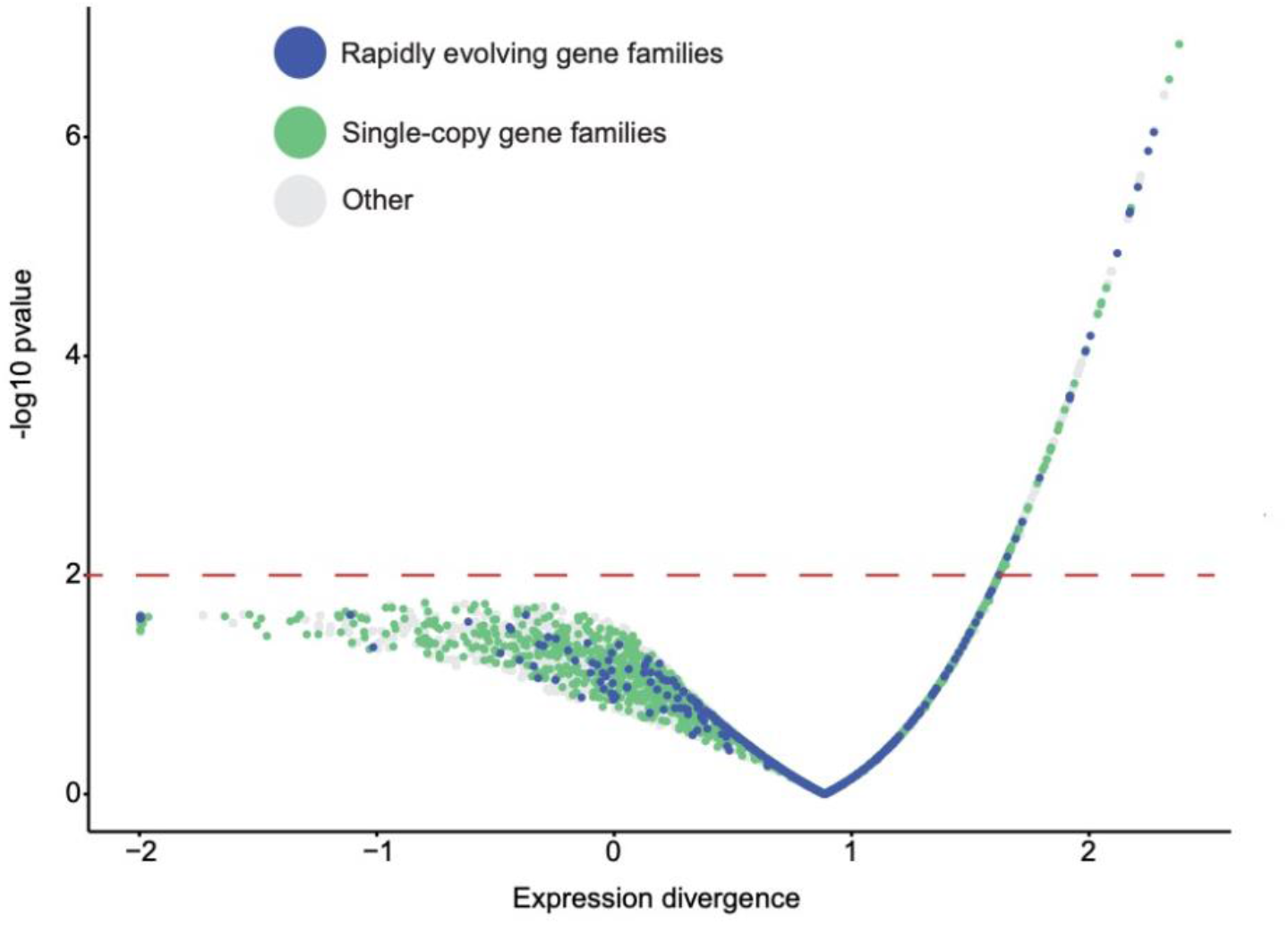
Gene family complexity is associated with species-level expression shifts. The rapidly evolving gene families have elevated expression divergence compared to single copy gene families (Odds ratio 1.93, p< 0.05) fisher’s exact test. The rapidly evolving gene families are in blue, the single copy gene families are in green and the other gene families are grey. Red line denotes significance (p<0.01).

In order to investigate if the abundance of immune related gene families which are rapidly evolving are translated into expression divergence we compared the 36 rapidly evolving immune gene families (Figure 4a,b) to the remaining 272 rapidly evolving gene families and the 1629 single copy gene families (Figure 4a,b). The immune related gene families have the highest expression divergence, followed by the rapidly evolving gene families, while single-copy gene families possess the lowest expression divergence, where all differences are significant (p<0.05) following a Tukeys’ HSD (figure 4a,b).

**Figure 4:**
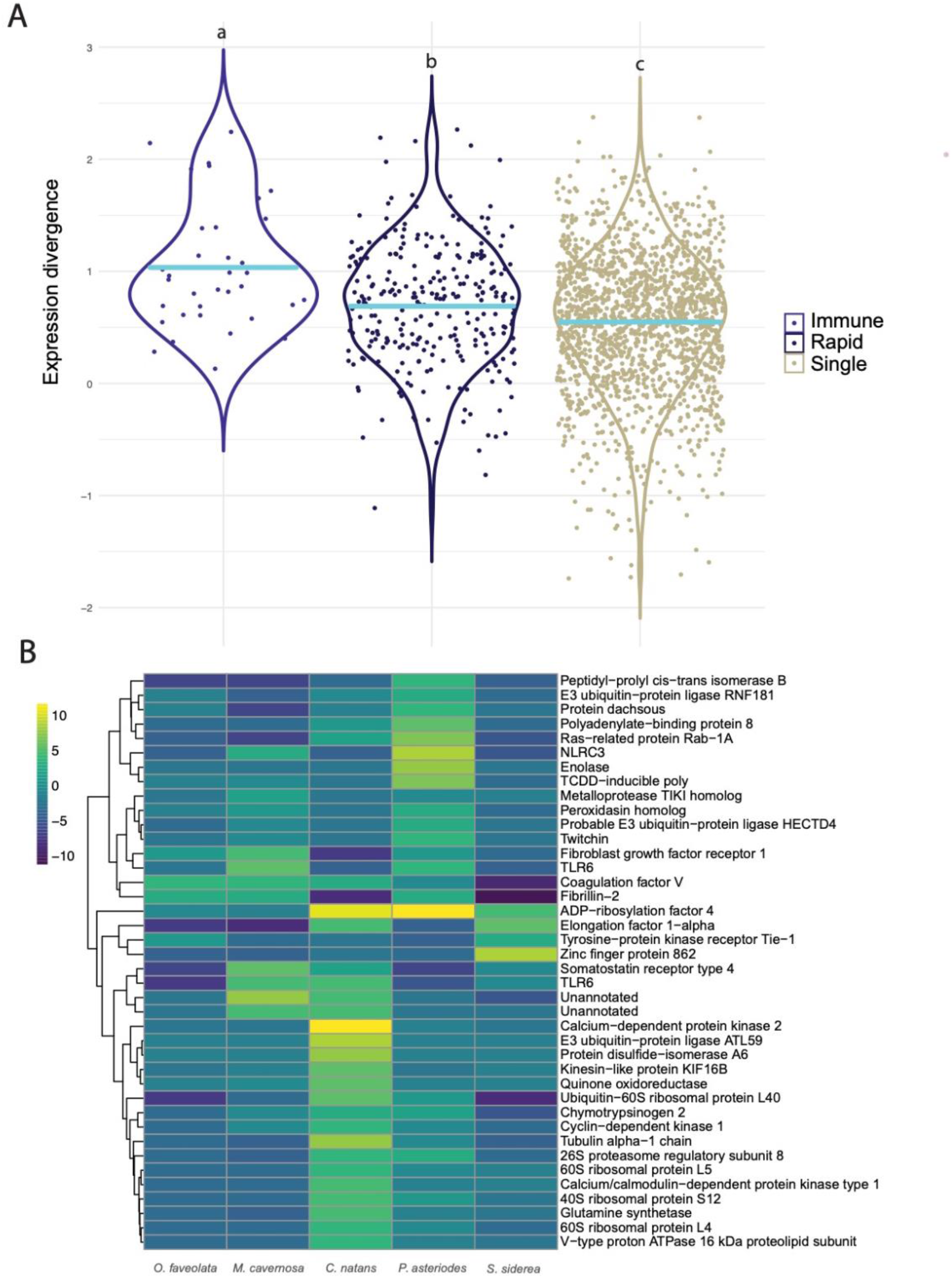
Rapid evolution of immune genes. A) The expression divergence is shown for rapidly evolving immune genes, rapidly evolving gene families, and single copy gene families. Group mean is represented by the blue line and all comparisons are significant after anova with Tukeys HSD multiple comparison test (p<0.01). B) Heatmap showing the predicted change in copy number for the rapidly evolving immune gene families compared to the last common ancestor. Gene names were given based upon the lowest Uniprot e-value for all of the contigs in the gene family. Gene families with a change of more than 15 copies in a species are shown at 15+.

## Discussion

Our investigation on the evolution of transcriptional complexity and expression divergence among five species of common Caribbean coral reveals several important findings. First, there are very few single copy orthologs between species indicating that variation in transcriptional complexity of gene families is widespread. Second, the genes with evolved changes in complexity occur in functionally related gene families affecting a number of biological processes, with immune and metabolism related gene families particularly well represented. Finally, transcriptional complexity of gene families promotes gene expression divergence. These results highlight the widespread divergence of gene repertoires as well as the role of transcriptional complexity as a mechanism of species-level adaptation in Caribbean corals.

In addition to demonstrating the prevalence of variation in gene repertoires among Caribbean coral, our study also demonstrates the feasibility of quantifying the evolution of gene family complexity in organisms which only have available transcriptomic resources. Fragmented and error-prone assemblies are expected to lead to inflated estimates of gene turnover. However, our calculated rate of gene family turnover approximates the observed rate among species of *Drosophila* (Han et al., 2013) and is lower than a study which utilized whole genomes from six species of mammals (Hahn, Demuth, & Han, 2007). This demonstrates that by employing stringent filtering approaches, errors common in transcriptome assemblies can be reduced to a level which allows investigation of transcriptional complexity evolution. Thus, increasing the accessibility of investigating this process in species currently lacking genome assemblies. In supports of this, our filtering approach yielded assemblies with a comparable number of predicted genes and BUSCO scores to existing whole-genome assemblies for corals (Shinzato et al., 2020, 2011; Shumaker et al., 2019; Voolstra et al., 2017). Additionally, assembly quality metrics were not associated with the number of rapid evolving gene families, further supporting the efficacy of our filtering approaches. Thus, this framework allowed us to expand our study to include ecologically important species currently lacking genome resources and provides a template for other investigations which seek to explore gene family evolution in non-model species in a cost-effective manner.

### Widespread Variation of Gene Repertoires

We identified 2009 single copy orthologs between the five species, meaning less than a third of the 6324 orthologous gene families present across the five species are conserved in a one-to-one relationship. This indicates that substantial variation in gene repertoires exist across these species. Interestingly, we observe a slow rate of gene turnover which may be a reflection of the long generation times and low mutation rates seen in corals (López & Palumbi, 2020). This means that the observed gene repertoire divergence is due to a slow build-up of variation across deep divergence times, rather than rapid rates of evolution. Thus, although reef-building corals within Scleractinia form a monophyletic group (Kayal et al., 2018), the ancient species splits (Pinzón C. et al., 2014) lead to substantial variation in the gene repertoires of each species. Given this variation, extrapolating gene centric findings across different species of corals needs to be performed with caution, as the one gene to one gene homologous relationship which is often assumed may be the exception rather that the rule. Thus, despite the slow rate of genome evolution, the deep divergence times between species of coral may mean that variation at the scale of genome architecture is more widespread than is currently recognized. This finding further highlights the need for studies of gene function and the production of additionally high-quality genomic resources from diverse genera of coral.

### Variation in Transcriptional Complexity as a Source for Adaptation

The variation in transcriptional complexity demonstrates two patterns of rapid evolution where three of the five species had an abundance of gene families with increased complexity, while the other two species had an abundance of gene families with decreased complexity. This shows that both putative gene gain and loss events were important forces in the evolutionary history of these organisms. In support of this all species had enriched terms in the gene families with increased transcriptional complexity, while two species had enrichments in the gene families with reduced complexity. This highlights transcriptional complexity as a source of adaptation to the complex and multi-faceted selective regimes these species have evolved under. Gene duplication as well as gene loss has been shown to facilitate adaptation in a variety of systems (Deng et al., 2010; Hazzouri et al., 2020; Swann, Holland, Petersen, Pietsch, & Boehm, 2020; Zhang, Zhang, & Rosenberg, 2002), and thus likely has contributed to the evolutionary history of these coral species.

The enriched terms span numerous biological processes both within and across species, however, some enriched terms do qualitatively correspond to host biology. For example, the species which can attain the largest colony diameter *O. faveolata* (Madin et al., 2016), has increased complexity of calcium ion transport genes which could relate to calcification and colony growth. Additionally, *M. cavernosa* has enrichments for the terms “response to food” and “digestion” among the gene families with increased complexity. *M. cavernosa* also has large polyps and a greater capacity for prey capture and heterotrophy (Houlbrèque & Ferrier-Pagès, 2009) demonstrating a potential link between species biology and the rapidly evolving gene families. Overall, this highlights that gene family evolution may be linked to ecologically and evolutionarily important traits.

### Gene family complexity promotes expression divergence

Rapidly evolving gene families are enriched for species-level expression shifts, demonstrating a link between expression divergence and transcriptional complexity. This pattern supports previous work showing that copy number influences gene dosage (Jiang & Assis, 2019) and can relax purifying selection promoting the processes of neofunctionalization and subfunctionalization (Zhang, 2003). Notably, gene families with significant variation in complexity have an overall higher level of expression divergence than single copy gene families, indicating that transcriptional complexity of a gene family may be a genetic pre-requisite to promote adaptive expression shifts (Wagner, 1994). Gene expression divergence can serve to promote adaptation (Rohlfs & Nielsen, 2015), and the link between expression divergence and transcriptional complexity further supports a role for this process as a source of adaptation. Given this data, multi-species comparisons should integrate gene family complexity given its influence on expression dynamics.

### Rapid evolution of Immunity

Immune genes are rapidly evolving in four of the five species investigated and demonstrate high levels of expression divergence. This reflects findings from other systems which have characterized the rapid evolution and subsequent change in copy number of immune genes (Evans et al., 2006; Sackton, Lazzaro, & Clark, 2017; Sackton et al., 2007; Waterhouse et al., 2007). This common pattern is thought to be due to the propensity of immune genes to engage in evolutionary arms races through their ability to mediate interactions with other organisms (Lazzaro & Clark, 2012). This process is likely important in reef-building corals that live in a microbe rich environment (van Oppen & Blackall, 2019). The immune system of corals functions to both regulate commensal microbes (beneficial bacteria and the algal *Symbiodiniaceae*) as well as defend against pathogenic ones (Kvennefors et al., 2010; Mansfield & Gilmore, 2019; Mydlarz, Jones, & Harvell, 2006; Wu et al., 2019). Thus, the rapid evolution of immune genes indicates that these interactions may have played an important role in shaping the evolutionary history of these species. Other investigations have found similar trends including phylosymbiosis between coral species and bacterial communities on the great barrier reef (Pollock et al., 2018), as well as highly stable host microbe associations in the Red sea (Ziegler et al., 2019). Overall, this indicates that interactions with microbes may broadly play a role in shaping the evolution of corals.

Of particular interest is the interaction formed with Symbiodiniaceae as this relationship requires the modulation of host immune processes (Mansfield & Gilmore, 2019; Matthews et al., 2017; Neubauer et al., 2017) in order to receive a supply of translocated carbohydrates from their symbionts. Different species of coral are known to preferentially associated with one or a few species of *Symbiodiniaceae* (Thornhill, Fitt, & Schmidt, 2006), although the mechanism by which hosts distinguish between these different symbionts is unknown. The rapid evolution of immune genes by the host may be one mechanism to facilitate this relationship through neofunctionalization of immune genes, however this hypothesis would need to be functionally confirmed. Different species of Symbiodiniaceae also differ in their metabolic profiles and amount of nutrients translocated to the host (Matthews et al., 2017; Wall, Kaluhiokalani, Popp, Donahue, & Gates, 2020). These differences may be reflected in host metabolic strategy and subsequent complexity of metabolic gene families. For example, both *O. faveolata* and *P. astreoides* have increased complexity of carbohydrate metabolism genes, *C. natans* and *S. siderea* have increased complexity of genes involved in amino acid metabolism, while *M. cavernosa* has increased complexity of lipid metabolism genes. These differences could potentially reflect an evolved mechanism to match host nutritional strategy with that of its preferred symbiotic partner. Considering the relationship between nutrient transfer and trophic strategy across different host-symbiont pairings is likely a fruitful area of investigation.

In addition to modulating the relationship with beneficial microorganisms, corals must also defend themselves against disease-causing microbes. This is particularly pressing in the Caribbean as disease outbreaks affecting reef-building corals have re-shaped communities (Aronson & Precht, 2001; Gardner, 2003; Miller et al., 2009; Mydlarz et al., 2006) and continue to be among the most pressing selective forces these organisms face (Vega Thurber et al., 2020). In response to disease, corals utilize a complex innate immune system involving pattern recognition receptors including Toll-like receptors (TLRs) (Poole & Weis, 2014; L. M. Williams et al., 2018) and NOD-like receptors (NLRs) (Dimos et al., 2019; Hamada et al., 2013) to sense threats and initiate immune responses. Interestingly, we observe that *M. cavernosa*, *C. natans* and *P. astreoides* have increased complexity of these immune recognition gene families, while *O. faveolata* has decreased complexity within these gene families. This variation in complexity of immune genes may influence disease-susceptibility. *O. faveolata* is highly susceptible to several coral diseases whereas *M. cavernosa* and *P. astreoides*, who have increased immune gene complexity, are considered more resilient to disease (Aeby et al., 2019; Smith et al., 2013; L. Williams, Smith, Burge, & Brandt, 2020). In support of this, species-level variation in immune activity is widespread in Caribbean corals (Palmer et al., 2011; Pinzón C. et al., 2014; Rosales, Clark, Huebner, Ruzicka, & Muller, 2020), which may be the result of the variation of immune gene repertoires we observe. However, *C. natans* is also considered to be susceptible to disease (Aeby et al., 2019; Sutherland, Porter, & Torres, 2004) and demonstrates increased complexity of some immune genes. Thus, the relationship between more complex immune gene repertoires and resistance to disease may not always lead to real-world immunocompetence.

Together these data indicate that the immune system is a major target of evolution in Caribbean coral. As coral diseases are currently among the strongest selective forces acting on Caribbean reefs (Vega Thurber et al., 2020) understanding the differing immune repertoires each species possesses and how this relates to disease resilience may be an important determinant of which corals will persist on the reef. Indeed, in the Caribbean, coral species which have historically dominated stable forereef environments such as *O. faveolata* have been declining in abundance (Gardner, 2003) while corals such as *P. astreoides* that have traditionally been found in the variable environment of the reef edge are rising in abundance (Green, Edmunds, & Carpenter, 2008; McWilliam, Pratchett, Hoogenboom, & Hughes, 2020) with disease playing a major role in these shifts.

## Conclusion

We found that putative changes in gene copy number are widespread in reef-building corals and that these changes influence both gene family complexity and expression. This finding supports mounting evidence that gene duplication serves as a mechanism of species-level adaptation in natural systems. Additionally, we show the rapid evolution of immune genes in coral, potentially owing to the complex nature of their associations with microorganisms. Overall, these findings have important implications from both an evolutionary and ecological perspective as they highlight previously unrecognized variation at the level of gene repertoire composition and demonstrates how this may impact coral biology and ecology. As selective pressures in the ocean change due to anthropogenic impacts and species compositions are reshaped as a result, it will become increasingly important to understand the adaptative capacity each species possesses. Therefore, further work towards linking variation in gene copy number to adaptive capacity may help predict which species will be able to adapt and which are likely to face extirpation.

## Author Contributions

### Data accessibility

All raw reads from this project are available on the NCBI short read archive under Bioproject number: PRJNA723585. All code and data needed to recreate the major findings from this paper are publicly available: https://github.com/braddimo/Gene-expansion.

### Author Contributions

B.D., M.E. and J.D. designed and preformed research. L.M. and M.E. assisted with writing. B.D. and N.M. performed lab work and constructed assemblies. M.B. assisted with sample collection.

## Acknowledgements

We would like to thank Marilyn Brandt and the student of the Center for Marine and Environmental Studies, University of the Virgin Islands for their help with sample collecting as well as the high performance computing center at the University of Texas at Arlington.

## References

Aeby, G. S., Ushijima, B., Campbell, J. E., Jones, S., Williams, G. J., Meyer, J. L., … Paul, V. J. (2019). Pathogenesis of a Tissue Loss Disease Affecting Multiple Species of Corals Along the Florida Reef Tract. Frontiers in Marine Science, 6, 678. doi: 10.3389/fmars.2019.00678

Alvarez-Filip, L., Dulvy, N. K., Gill, J. A., Côté, I. M., & Watkinson, A. R. (2009). Flattening of Caribbean coral reefs: Region-wide declines in architectural complexity. Proceedings of the Royal Society B: Biological Sciences, 276(1669), 3019–3025. doi: 10.1098/rspb.2009.0339

Aranda, M., Li, Y., Liew, Y. J., Baumgarten, S., Simakov, O., Wilson, M. C., … Voolstra, C. R. (2016). Genomes of coral dinoflagellate symbionts highlight evolutionary adaptations conducive to a symbiotic lifestyle. Scientific Reports, 6(1), 39734. doi: 10.1038/srep39734

Aronson, R. B., & Precht, W. F. (2001). White-band disease and the changing face of Caribbean coral reefs. In J. W. Porter (Ed.), The Ecology and Etiology of Newly Emerging Marine Diseases (pp. 25–38). Dordrecht: Springer Netherlands. doi: 10.1007/978-94-017-3284-0_2

Baker, A. C., Glynn, P. W., & Riegl, B. (2008). Climate change and coral reef bleaching: An ecological assessment of long-term impacts, recovery trends and future outlook. Estuarine, Coastal and Shelf Science, 80(4), 435–471. doi: 10.1016/j.ecss.2008.09.003

Bay, R. A., & Palumbi, S. R. (2014). Multilocus Adaptation Associated with Heat Resistance in Reef-Building Corals. Current Biology, 24(24), 2952–2956. doi: 10.1016/j.cub.2014.10.044

Bay, R. A., & Palumbi, S. R. (2015). Rapid Acclimation Ability Mediated by Transcriptome Changes in Reef-Building Corals. Genome Biology and Evolution, 7(6), 1602–1612. doi: 10.1093/gbe/evv085

Bolger, A. M., Lohse, M., & Usadel, B. (2014). Trimmomatic: A flexible trimmer for Illumina sequence data. Bioinformatics, 30(15), 2114–2120. doi: 10.1093/bioinformatics/btu170

Brown, C. J., Todd, K. M., & Rosenzweig, R. F. (1998). Multiple duplications of yeast hexose transport genes in response to selection in a glucose-limited environment. Molecular Biology and Evolution, 15(8), 931–942. doi: 10.1093/oxfordjournals.molbev.a026009

Chain, F. J. J., Feulner, P. G. D., Panchal, M., Eizaguirre, C., Samonte, I. E., Kalbe, M., … Reusch, T. B. H. (2014). Extensive Copy-Number Variation of Young Genes across Stickleback Populations. PLoS Genetics, 10(12), e1004830. doi: 10.1371/journal.pgen.1004830

Davies, S. W., Treml, E. A., Kenkel, C. D., & Matz, M. V. (2015). Exploring the role of Micronesian islands in the maintenance of coral genetic diversity in the Pacific Ocean. Molecular Ecology, 24(1), 70–82. doi: 10.1111/mec.13005

Davies, Sarah W., Marchetti, A., Ries, J. B., & Castillo, K. D. (2016). Thermal and pCO2 Stress Elicit Divergent Transcriptomic Responses in a Resilient Coral. Frontiers in Marine Science, 3. doi: 10.3389/fmars.2016.00112

De Smet, R., Sabaghian, E., Li, Z., Saeys, Y., & Van de Peer, Y. (2017). Coordinated Functional Divergence of Genes after Genome Duplication in *Arabidopsis thaliana*. The Plant Cell, 29(11), 2786–2800. doi: 10.1105/tpc.17.00531

Deng, C., Cheng, C.-H. C., Ye, H., He, X., & Chen, L. (2010). Evolution of an antifreeze protein by neofunctionalization under escape from adaptive conflict. Proceedings of the National Academy of Sciences, 107(50), 21593–21598. doi: 10.1073/pnas.1007883107

Dimos, B. A., Butler, C. C., Ricci, C. A., MacKnight, N. J., & Mydlarz, L. D. (2019). Responding to Threats Both Foreign and Domestic: NOD-Like Receptors in Corals. Integrative and Comparative Biology, 59(4), 819–829. doi: 10.1093/icb/icz111

Emms, David M., & Kelly, S. (2015). OrthoFinder: Solving fundamental biases in whole genome comparisons dramatically improves orthogroup inference accuracy. Genome Biology, 16(1), 157. doi: 10.1186/s13059-015-0721-2

Emms, David M., & Kelly, S. (2019). OrthoFinder: Phylogenetic orthology inference for comparative genomics. Genome Biology, 20(1), 238. doi: 10.1186/s13059-019-1832-y

Emms, D.M., & Kelly, S. (2018). *STAG: Species Tree Inference from All Genes* [Preprint]. Evolutionary Biology. doi: 10.1101/267914

Evans, J. D., Aronstein, K., Chen, Y. P., Hetru, C., Imler, J.-L., Jiang, H., … Hultmark, D. (2006). Immune pathways and defence mechanisms in honey bees Apis mellifera. Insect Molecular Biology, 15(5), 645–656. doi: 10.1111/j.1365-2583.2006.00682.x

Force, A., Lynch, M., Pickett, F. B., Amores, A., Yan, Y. L., & Postlethwait, J. (1999). Preservation of duplicate genes by complementary, degenerative mutations. Genetics, 151(4), 1531– 1545.

Fraser, H. B. (2011). Genome-wide approaches to the study of adaptive gene expression evolution: Systematic studies of evolutionary adaptations involving gene expression will allow many fundamental questions in evolutionary biology to be addressed. BioEssays, 33(6), 469–477. doi: 10.1002/bies.201000094

Fuller, Z. L., Mocellin, V. J. L., Morris, L. A., Cantin, N., Shepherd, J., Sarre, L., … Przeworski, M. (2020). Population genetics of the coral Acropora millepora: Toward genomic prediction of bleaching. Science (New York, N.Y.), 369(6501). doi: 10.1126/science.aba4674

Gardner, T. A. (2003). Long-Term Region-Wide Declines in Caribbean Corals. Science, 301(5635), 958–960. doi: 10.1126/science.1086050

Gilad, Y., Oshlack, A., & Rifkin, S. A. (2006). Natural selection on gene expression. Trends in Genetics, 22(8), 456–461. doi: 10.1016/j.tig.2006.06.002

Grabherr, M. G., Haas, B. J., Yassour, M., Levin, J. Z., Thompson, D. A., Amit, I., … Regev, A. (2011). Full-length transcriptome assembly from RNA-Seq data without a reference genome. Nature Biotechnology, 29(7), 644–652. doi: 10.1038/nbt.1883

Green, D., Edmunds, P., & Carpenter, R. (2008). Increasing relative abundance of Porites astreoides on Caribbean reefs mediated by an overall decline in coral cover. Marine Ecology Progress Series, 359, 1–10. doi: 10.3354/meps07454

Haas, B. J., Papanicolaou, A., Yassour, M., Grabherr, M., Blood, P. D., Bowden, J., … Regev, A. (2013). De novo transcript sequence reconstruction from RNA-seq using the Trinity platform for reference generation and analysis. Nature Protocols, 8(8), 1494–1512. doi: 10.1038/nprot.2013.084

Hahn, M. W., Demuth, J. P., & Han, S.-G. (2007). Accelerated Rate of Gene Gain and Loss in Primates. Genetics, 177(3), 1941–1949. doi: 10.1534/genetics.107.080077

Hamada, M., Shoguchi, E., Shinzato, C., Kawashima, T., Miller, D. J., & Satoh, N. (2013). The Complex NOD-Like Receptor Repertoire of the Coral Acropora digitifera Includes Novel Domain Combinations. Molecular Biology and Evolution, 30(1), 167–176. doi: 10.1093/molbev/mss213

Han, M. V., Thomas, G. W. C., Lugo-Martinez, J., & Hahn, M. W. (2013). Estimating Gene Gain and Loss Rates in the Presence of Error in Genome Assembly and Annotation Using CAFE 3. Molecular Biology and Evolution, 30(8), 1987–1997. doi: 10.1093/molbev/mst100

Hazzouri, K. M., Sudalaimuthuasari, N., Kundu, B., Nelson, D., Al-Deeb, M. A., Le Mansour, A., … Amiri, K. M. A. (2020). The genome of pest Rhynchophorus ferrugineus reveals gene families important at the plant-beetle interface. Communications Biology, 3(1), 323. doi: 10.1038/s42003-020-1060-8

Hirase, S., Ozaki, H., & Iwasaki, W. (2014). Parallel selection on gene copy number variations through evolution of three-spined stickleback genomes. BMC Genomics, 15(1), 735. doi: 10.1186/1471-2164-15-735

Houlbrèque, F., & Ferrier-Pagès, C. (2009). Heterotrophy in Tropical Scleractinian Corals. Biological Reviews, 84(1), 1–17. doi: 10.1111/j.1469-185X.2008.00058.x

Huang, K. M., & Chain, F. J. J. (2021). Copy number variations and young duplicate genes have high methylation levels in sticklebacks. Evolution, 75(3), 706–718. doi: 10.1111/evo.14184

Huang, Ying, Niu, B., Gao, Y., Fu, L., & Li, W. (2010). CD-HIT Suite: A web server for clustering and comparing biological sequences. Bioinformatics, 26(5), 680–682. doi: 10.1093/bioinformatics/btq003

Huang, Yun, Feulner, P. G. D., Eizaguirre, C., Lenz, T. L., Bornberg-Bauer, E., Milinski, M., … Chain, F. J. J. (2019). Genome-Wide Genotype-Expression Relationships Reveal Both Copy Number and Single Nucleotide Differentiation Contribute to Differential Gene Expression between Stickleback Ecotypes. Genome Biology and Evolution, 11(8), 2344– 2359. doi: 10.1093/gbe/evz148

Huerta-Cepas, J., Forslund, K., Coelho, L. P., Szklarczyk, D., Jensen, L. J., von Mering, C., & Bork, P. (2017). Fast Genome-Wide Functional Annotation through Orthology Assignment by eggNOG-Mapper. Molecular Biology and Evolution, 34(8), 2115–2122. doi: 10.1093/molbev/msx148

Huerta-Cepas, J., Szklarczyk, D., Heller, D., Hernández-Plaza, A., Forslund, S. K., Cook, H., … Bork, P. (2019). eggNOG 5.0: A hierarchical, functionally and phylogenetically annotated orthology resource based on 5090 organisms and 2502 viruses. Nucleic Acids Research, 47(D1), D309–D314. doi: 10.1093/nar/gky1085

Jiang, X., & Assis, R. (2019). Rapid functional divergence after small-scale gene duplication in grasses. BMC Evolutionary Biology, 19(1), 97. doi: 10.1186/s12862-019-1415-2

Kayal, E., Bentlage, B., Sabrina Pankey, M., Ohdera, A. H., Medina, M., Plachetzki, D. C., … Ryan, J. F. (2018). Phylogenomics provides a robust topology of the major cnidarian lineages and insights on the origins of key organismal traits. BMC Evolutionary Biology, 18(1), 68. doi: 10.1186/s12862-018-1142-0

Kenkel, C. D., & Matz, M. V. (2017). Gene expression plasticity as a mechanism of coral adaptation to a variable environment. Nature Ecology & Evolution, 1(1), 0014. doi: 10.1038/s41559-016-0014

Kim, D., Pertea, G., Trapnell, C., Pimentel, H., Kelley, R., & Salzberg, S. L. (2013). TopHat2: Accurate alignment of transcriptomes in the presence of insertions, deletions and gene fusions. Genome Biology, 14(4), R36. doi: 10.1186/gb-2013-14-4-r36

Kirk, N. L., Howells, E. J., Abrego, D., Burt, J. A., & Meyer, E. (2017). Genomic and transcriptomic signals of thermal tolerance in heat-tolerant corals (Platygyra daedalea) of the Arabian/Persian Gulf. doi: 10.1101/185579

Kondrashov, F. A. (2012). Gene duplication as a mechanism of genomic adaptation to a changing environment. Proceedings of the Royal Society B: Biological Sciences, 279(1749), 5048–5057. doi: 10.1098/rspb.2012.1108

Kvennefors, E. C. E., Leggat, W., Kerr, C. C., Ainsworth, T. D., Hoegh-Guldberg, O., & Barnes, A. C. (2010). Analysis of evolutionarily conserved innate immune components in coral links immunity and symbiosis. Developmental & Comparative Immunology, 34(11), 1219– 1229. doi: 10.1016/j.dci.2010.06.016

Lazzaro, B. P., & Clark, A. G. (2012). Rapid evolution of innate immune response genes. In R. S. Singh, J. Xu, & R. J. Kulathinal (Eds.), Rapidly Evolving Genes and Genetic Systems (pp. 203–210). Oxford University Press. doi: 10.1093/acprof:oso/9780199642274.003.0020

Liu, H., Stephens, T. G., González-Pech, R. A., Beltran, V. H., Lapeyre, B., Bongaerts, P., … Chan, C. X. (2018). Symbiodinium genomes reveal adaptive evolution of functions related to coral-dinoflagellate symbiosis. Communications Biology, 1(1), 95. doi: 10.1038/s42003-018-0098-3

López, E. H., & Palumbi, S. R. (2020). Somatic Mutations and Genome Stability Maintenance in Clonal Coral Colonies. Molecular Biology and Evolution, 37(3), 828–838. doi: 10.1093/molbev/msz270

Love, M. I., Huber, W., & Anders, S. (2014). Moderated estimation of fold change and dispersion for RNA-seq data with DESeq2. Genome Biology, 15(12). doi: 10.1186/s13059-014-0550-8

Madin, J. S., Anderson, K. D., Andreasen, M. H., Bridge, T. C. L., Cairns, S. D., Connolly, S. R., … Baird, A. H. (2016). The Coral Trait Database, a curated database of trait information for coral species from the global oceans. Scientific Data, 3(1), 160017. doi: 10.1038/sdata.2016.17

Mansfield, K. M., & Gilmore, T. D. (2019). Innate immunity and cnidarian-Symbiodiniaceae mutualism. Developmental & Comparative Immunology, 90, 199–209. doi: 10.1016/j.dci.2018.09.020

Matthews, J. L., Crowder, C. M., Oakley, C. A., Lutz, A., Roessner, U., Meyer, E., … Davy, S. K. (2017). Optimal nutrient exchange and immune responses operate in partner specificity in the cnidarian-dinoflagellate symbiosis. Proceedings of the National Academy of Sciences, 114(50), 13194–13199. doi: 10.1073/pnas.1710733114

Matz, M. V., Treml, E. A., Aglyamova, G. V., & Bay, L. K. (2018). Potential and limits for rapid genetic adaptation to warming in a Great Barrier Reef coral. PLOS Genetics, 14(4), e1007220. doi: 10.1371/journal.pgen.1007220

Maynard, J., van Hooidonk, R., Eakin, C. M., Puotinen, M., Garren, M., Williams, G., … Harvell, C. C. (2015). Projections of climate conditions that increase coral disease susceptibility and pathogen abundance and virulence. Nature Climate Change, 5(7), 688–694. doi: 10.1038/nclimate2625

McWilliam, M., Pratchett, M. S., Hoogenboom, M. O., & Hughes, T. P. (2020). Deficits in functional trait diversity following recovery on coral reefs. Proceedings of the Royal Society B: Biological Sciences, 287(1918), 20192628. doi: 10.1098/rspb.2019.2628

Meyers, B. C., Vu, T. H., Tej, S. S., Ghazal, H., Matvienko, M., Agrawal, V., … Haudenschild, C. D. (2004). Analysis of the transcriptional complexity of Arabidopsis thaliana by massively parallel signature sequencing. Nature Biotechnology, 22(8), 1006–1011. doi: 10.1038/nbt992

Miller, J., Muller, E., Rogers, C., Waara, R., Atkinson, A., Whelan, K. R. T., … Witcher, B. (2009). Coral disease following massive bleaching in 2005 causes 60% decline in coral cover on reefs in the US Virgin Islands. Coral Reefs, 28(4), 925–937. doi: 10.1007/s00338-009-0531-7

Monks, S. A., Leonardson, A., Zhu, H., Cundiff, P., Pietrusiak, P., Edwards, S., … Schadt, E. E. (2004). Genetic Inheritance of Gene Expression in Human Cell Lines. The American Journal of Human Genetics, 75(6), 1094–1105. doi: 10.1086/426461

Moya, A., Huisman, L., Ball, E. E., Hayward, D. C., Grasso, L. C., Chua, C. M., … Miller, D. J. (2012). Whole Transcriptome Analysis of the Coral *Acropora millepora* Reveals Complex Responses to CO _2_ -driven Acidification during the Initiation of Calcification: TRANSCRIPTOMIC RESPONSE OF CORALS TO ACIDIFICATION. Molecular Ecology, 21(10), 2440–2454. doi: 10.1111/j.1365-294X.2012.05554.x

Mydlarz, L. D., Jones, L. E., & Harvell, C. D. (2006). Innate Immunity, Environmental Drivers, and Disease Ecology of Marine and Freshwater Invertebrates. Annual Review of Ecology, Evolution, and Systematics, 37(1), 251–288. doi: 10.1146/annurev.ecolsys.37.091305.110103

Neubauer, E.-F., Poole, A. Z., Neubauer, P., Detournay, O., Tan, K., Davy, S. K., & Weis, V. M. (2017). A diverse host thrombospondin-type-1 repeat protein repertoire promotes symbiont colonization during establishment of cnidarian-dinoflagellate symbiosis. ELife, 6, e24494. doi: 10.7554/eLife.24494

Ohno, Susumu. (1970). Evolution by Gene Duplication. New York: Springer.

Palmer, C. V., McGinty, E. S., Cummings, D. J., Smith, S. M., Bartels, E., & Mydlarz, L. D. (2011). Patterns of coral ecological immunology: Variation in the responses of Caribbean corals to elevated temperature and a pathogen elicitor. Journal of Experimental Biology, 214(24), 4240–4249. doi: 10.1242/jeb.061267

Parkinson, J. E., Baumgarten, S., Michell, C. T., Baums, I. B., LaJeunesse, T. C., & Voolstra, C. R. (2016). Gene Expression Variation Resolves Species and Individual Strains among Coral-Associated Dinoflagellates within the Genus *Symbiodinium*. Genome Biology and Evolution, 8(3), 665–680. doi: 10.1093/gbe/evw019

Patro, R., Duggal, G., Love, M. I., Irizarry, R. A., & Kingsford, C. (2017). Salmon provides fast and bias-aware quantification of transcript expression. Nature Methods, 14(4), 417–419. doi: 10.1038/nmeth.4197

Pinzón C., J. H., Beach-Letendre, J., Weil, E., & Mydlarz, L. D. (2014). Relationship between Phylogeny and Immunity Suggests Older Caribbean Coral Lineages Are More Resistant to Disease. PLoS ONE, 9(8), e104787. doi: 10.1371/journal.pone.0104787

Pollock, F. J., McMinds, R., Smith, S., Bourne, D. G., Willis, B. L., Medina, M., … Zaneveld, J. R. (2018). Coral-associated bacteria demonstrate phylosymbiosis and cophylogeny. Nature Communications, 9(1), 4921. doi: 10.1038/s41467-018-07275-x

Poole, A. Z., & Weis, V. M. (2014). TIR-domain-containing protein repertoire of nine anthozoan species reveals coral–specific expansions and uncharacterized proteins. Developmental & Comparative Immunology, 46(2), 480–488. doi: 10.1016/j.dci.2014.06.002

Prada, C., Hanna, B., Budd, A. F., Woodley, C. M., Schmutz, J., Grimwood, J., … Medina, M. (2016). Empty Niches after Extinctions Increase Population Sizes of Modern Corals. Current Biology: CB, 26(23), 3190–3194. doi: 10.1016/j.cub.2016.09.039

Precht, W. F., Gintert, B. E., Robbart, M. L., Fura, R., & van Woesik, R. (2016). Unprecedented Disease-Related Coral Mortality in Southeastern Florida. Scientific Reports, 6(1), 31374. doi: 10.1038/srep31374

Price, A. L., Helgason, A., Thorleifsson, G., McCarroll, S. A., Kong, A., & Stefansson, K. (2011). Single-Tissue and Cross-Tissue Heritability of Gene Expression Via Identity-by-Descent in Related or Unrelated Individuals. PLoS Genetics, 7(2), e1001317. doi: 10.1371/journal.pgen.1001317

Prunier, J., Caron, S., & MacKay, J. (2017). CNVs into the wild: Screening the genomes of conifer trees (Picea spp.) reveals fewer gene copy number variations in hybrids and links to adaptation. BMC Genomics, 18(1), 97. doi: 10.1186/s12864-016-3458-8

Putnam, H. M., Barott, K. L., Ainsworth, T. D., & Gates, R. D. (2017). The Vulnerability and Resilience of Reef-Building Corals. Current Biology, 27(11), R528–R540. doi: 10.1016/j.cub.2017.04.047

R Core Team (2020). R: A language and environment for statistical computing. R Foundation for Statistical Computing, Vienna, Austria. URL https://www.R-project.org/. (n.d.).

Revell, L. J. (2012). phytools: An R package for phylogenetic comparative biology (and other things): phytools: R package. Methods in Ecology and Evolution, 3(2), 217–223. doi: 10.1111/j.2041-210X.2011.00169.x

Rohlfs, R. V., & Nielsen, R. (2015). Phylogenetic ANOVA: The Expression Variance and Evolution Model for Quantitative Trait Evolution. Systematic Biology, 64(5), 695–708. doi: 10.1093/sysbio/syv042

Rosales, S. M., Clark, A. S., Huebner, L. K., Ruzicka, R. R., & Muller, E. M. (2020). Rhodobacterales and Rhizobiales Are Associated With Stony Coral Tissue Loss Disease and Its Suspected Sources of Transmission. Frontiers in Microbiology, 11, 681. doi: 10.3389/fmicb.2020.00681

Sackton, T. B., Lazzaro, B. P., & Clark, A. G. (2017). Rapid expansion of immune-related gene families in the house fly, *Musca domestica*. *Molecular Biology and Evolution*, msw285. doi: 10.1093/molbev/msw285

Sackton, T. B., Lazzaro, B. P., Schlenke, T. A., Evans, J. D., Hultmark, D., & Clark, A. G. (2007). Dynamic evolution of the innate immune system in Drosophila. Nature Genetics, 39(12), 1461–1468. doi: 10.1038/ng.2007.60

Schuster-Böckler, B., Conrad, D., & Bateman, A. (2010). Dosage Sensitivity Shapes the Evolution of Copy-Number Varied Regions. PLoS ONE, 5(3), e9474. doi: 10.1371/journal.pone.0009474

Shinzato, C., Khalturin, K., Inoue, J., Zayasu, Y., Kanda, M., Kawamitsu, M., … Satoh, N. (2020). Eighteen Coral Genomes Reveal the Evolutionary Origin of Acropora Strategies to Accommodate Environmental Changes. Molecular Biology and Evolution, msaa216. doi: 10.1093/molbev/msaa216

Shinzato, C., Shoguchi, E., Kawashima, T., Hamada, M., Hisata, K., Tanaka, M., … Satoh, N. (2011). Using the Acropora digitifera genome to understand coral responses to environmental change. Nature, 476(7360), 320–323. doi: 10.1038/nature10249

Shoguchi, E., Beedessee, G., Hisata, K., Tada, I., Narisoko, H., Satoh, N., … Shinzato, C. (2021). A New Dinoflagellate Genome Illuminates a Conserved Gene Cluster Involved in Sunscreen Biosynthesis. Genome Biology and Evolution, 13(2), evaa235. doi: 10.1093/gbe/evaa235

Shumaker, A., Putnam, H. M., Qiu, H., Price, D. C., Zelzion, E., Harel, A., … Bhattacharya, D. (2019). Genome analysis of the rice coral Montipora capitata. Scientific Reports, 9(1), 2571. doi: 10.1038/s41598-019-39274-3

Simão, F. A., Waterhouse, R. M., Ioannidis, P., Kriventseva, E. V., & Zdobnov, E. M. (2015). BUSCO: Assessing genome assembly and annotation completeness with single-copy orthologs. Bioinformatics, 31(19), 3210–3212. doi: 10.1093/bioinformatics/btv351

Smith, T. B., Brandt, M. E., Calnan, J. M., Nemeth, R. S., Blondeau, J., Kadison, E., … Rothenberger, P. (2013). Convergent mortality responses of Caribbean coral species to seawater warming. Ecosphere, 4(7), art87. doi: 10.1890/ES13-00107.1

Soneson, C., Love, M. I., & Robinson, M. D. (2015). Differential analyses for RNA-seq: Transcript-level estimates improve gene-level inferences. F1000Research, 4, 1521. doi: 10.12688/f1000research.7563.1

Stuart-Smith, R. D., Brown, C. J., Ceccarelli, D. M., & Edgar, G. J. (2018). Ecosystem restructuring along the Great Barrier Reef following mass coral bleaching. Nature, 560(7716), 92–96. doi: 10.1038/s41586-018-0359-9

Sutherland, K., Porter, J., & Torres, C. (2004). Disease and immunity in Caribbean and Indo-Pacific zooxanthellate corals. Marine Ecology Progress Series, 266, 273–302. doi: 10.3354/meps266273

Swann, J. B., Holland, S. J., Petersen, M., Pietsch, T. W., & Boehm, T. (2020). The immunogenetics of sexual parasitism. Science, 369(6511), 1608–1615. doi: 10.1126/science.aaz9445

Tejada-Martinez, D., de Magalhães, J. P., & Opazo, J. C. (2021). Positive selection and gene duplications in tumour suppressor genes reveal clues about how cetaceans resist cancer. Proceedings of the Royal Society B: Biological Sciences, 288(1945), 20202592. doi: 10.1098/rspb.2020.2592

The UniProt Consortium, Bateman, A., Martin, M.-J., Orchard, S., Magrane, M., Agivetova, R., … Teodoro, D. (2021). UniProt: The universal protein knowledgebase in 2021. Nucleic Acids Research, 49(D1), D480–D489. doi: 10.1093/nar/gkaa1100

Thornhill, D. J., Fitt, W. K., & Schmidt, G. W. (2006). Highly stable symbioses among western Atlantic brooding corals. Coral Reefs, 25(4), 515–519. doi: 10.1007/s00338-006-0157-y

van de Water, J. A. J. M., Chaib De Mares, M., Dixon, G. B., Raina, J.-B., Willis, B. L., Bourne, D. G., & van Oppen, M. J. H. (2018). Antimicrobial and stress responses to increased temperature and bacterial pathogen challenge in the holobiont of a reef-building coral. Molecular Ecology, 27(4), 1065–1080. doi: 10.1111/mec.14489

van Oppen, M. J. H., & Blackall, L. L. (2019). Coral microbiome dynamics, functions and design in a changing world. Nature Reviews Microbiology, 17(9), 557–567. doi: 10.1038/s41579-019-0223-4

van Woesik, R., & Randall, C. J. (2017). Coral disease hotspots in the Caribbean. Ecosphere, 8(5), e01814. doi: 10.1002/ecs2.1814

Vega Thurber, R., Mydlarz, L. D., Brandt, M., Harvell, D., Weil, E., Raymundo, L., … Lamb, J. (2020). Deciphering Coral Disease Dynamics: Integrating Host, Microbiome, and the Changing Environment. Frontiers in Ecology and Evolution, 8, 575927. doi: 10.3389/fevo.2020.575927

Voolstra, C. R., Li, Y., Liew, Y. J., Baumgarten, S., Zoccola, D., Flot, J.-F., … Aranda, M. (2017). Comparative analysis of the genomes of Stylophora pistillata and Acropora digitifera provides evidence for extensive differences between species of corals. Scientific Reports, 7(1), 17583. doi: 10.1038/s41598-017-17484-x

Wagner, A. (1994). Evolution of gene networks by gene duplications: A mathematical model and its implications on genome organization. Proceedings of the National Academy of Sciences, 91(10), 4387–4391. doi: 10.1073/pnas.91.10.4387

Wall, C. B., Kaluhiokalani, M., Popp, B. N., Donahue, M. J., & Gates, R. D. (2020). Divergent symbiont communities determine the physiology and nutrition of a reef coral across a light-availability gradient. The ISME Journal, 14(4), 945–958. doi: 10.1038/s41396-019-0570-1

Walton, C. J., Hayes, N. K., & Gilliam, D. S. (2018). Impacts of a Regional, Multi-Year, Multi-Species Coral Disease Outbreak in Southeast Florida. Frontiers in Marine Science, 5, 323. doi: 10.3389/fmars.2018.00323

Waterhouse, R. M., Kriventseva, E. V., Meister, S., Xi, Z., Alvarez, K. S., Bartholomay, L. C., … Christophides, G. K. (2007). Evolutionary Dynamics of Immune-Related Genes and Pathways in Disease-Vector Mosquitoes. Science, 316(5832), 1738–1743. doi: 10.1126/science.1139862

Whitehead, A., & Crawford, D. L. (2006). Variation within and among species in gene expression: Raw material for evolution: REVIEW OF GENE EXPRESSION VARIATION. Molecular Ecology, 15(5), 1197–1211. doi: 10.1111/j.1365-294X.2006.02868.x

Williams, L. M., Fuess, L. E., Brennan, J. J., Mansfield, K. M., Salas-Rodriguez, E., Welsh, J., … Gilmore, T. D. (2018). A conserved Toll-like receptor-to-NF-κB signaling pathway in the endangered coral Orbicella faveolata. Developmental & Comparative Immunology, 79, 128–136. doi: 10.1016/j.dci.2017.10.016

Williams, L., Smith, T. B., Burge, C. A., & Brandt, M. E. (2020). Species-specific susceptibility to white plague disease in three common Caribbean corals. Coral Reefs, 39(1), 27–31. doi: 10.1007/s00338-019-01867-9

Wright, R. M., Aglyamova, G. V., Meyer, E., & Matz, M. V. (2015). Gene expression associated with white syndromes in a reef building coral, Acropora hyacinthus. BMC Genomics, 16(1). doi: 10.1186/s12864-015-1540-2

Wu, Y., Zhou, Z., Wang, J., Luo, J., Wang, L., & Zhang, Y. (2019). Temperature regulates the recognition activities of a galectin to pathogen and symbiont in the scleractinian coral Pocillopora damicornis. Developmental & Comparative Immunology, 96, 103–110. doi: 10.1016/j.dci.2019.03.003

Würschum, T., Boeven, P. H. G., Langer, S. M., Longin, C. F. H., & Leiser, W. L. (2015). Multiply to conquer: Copy number variations at Ppd-B1 and Vrn-A1 facilitate global adaptation in wheat. BMC Genetics, 16(1), 96. doi: 10.1186/s12863-015-0258-0

Zhang, J. (2003). Evolution by gene duplication: An update. Trends in Ecology & Evolution, 18(6), 292–298. doi: 10.1016/S0169-5347(03)00033-8

Zhang, J., Zhang, Y., & Rosenberg, H. F. (2002). Adaptive evolution of a duplicated pancreatic ribonuclease gene in a leaf-eating monkey. Nature Genetics, 30(4), 411–415. doi: 10.1038/ng852

Ziegler, M., Grupstra, C. G. B., Barreto, M. M., Eaton, M., BaOmar, J., Zubier, K., … Voolstra, C. R. (2019). Coral bacterial community structure responds to environmental change in a host-specific manner. Nature Communications, 10(1), 3092. doi: 10.1038/s41467-019-10969-5

